# Latent taste diversity revealed by a vertebrate-wide catalogue of T1R receptors

**DOI:** 10.1101/2023.04.08.532961

**Authors:** Hidenori Nishihara, Yasuka Toda, Tae Kuramoto, Kota Kamohara, Azusa Goto, Kyoko Hoshino, Shinji Okada, Shigehiro Kuraku, Masataka Okabe, Yoshiro Ishimaru

## Abstract

Taste is a vital chemical sense for feeding behavior. In mammals, the umami and sweet taste receptors are composed of three members of the taste receptor type 1 (T1R/TAS1R) family: T1R1, T1R2, and T1R3. Because their functional homologs exist in teleosts, only three *TAS1R* genes generated by gene duplication are believed to have been inherited from the common ancestor of bony vertebrates. Here, we report five previously uncharacterized *TAS1R* members in vertebrates, named *TAS1R4*, *5*, *6*, *7*, and *TAS1Rcf*, through a genome-wide survey of diverse taxa. For *TAS1R2* and *TAS1R3*, mammalian and teleost fish genes were found to be paralogous. Phylogenetic analysis suggests that the bony vertebrate ancestor had nine *TAS1Rs* due to multiple gene duplications, and some *TAS1R*s were lost independently in each lineage; ultimately, mammals and teleosts have retained only three *TAS1R*s, whereas other lineages have retained more *TAS1Rs*. Functional assays and expression analysis in non-teleost fishes suggest that the novel T1Rs form heterodimers in taste receptor cells and contribute to the recognition of a broad range of ligands such as essential amino acids, including branched-chain amino acids, which were not previously considered as T1R ligands. These results highlight an unexpected diversity of taste sensations in both modern and the ancestors of vertebrates. The complex evolution of the taste receptor family might have enabled vertebrates to adapt to diverse habitats on Earth.

## Introduction

Taste is one of the most important senses that govern the feeding behavior of animals. It is widely accepted that mammals have five basic tastes: umami (savory), sweet, bitter, salty, and sour ^1, 2^. Taste receptor type 1 (T1R, encoded by *TAS1R*), a G protein–coupled receptor family, consists of three members, namely T1R1, T1R2, and T1R3, which are encoded by the genes *TAS1R1*, *TAS1R2*, and *TAS1R3*, respectively, and act as umami or sweet receptors ^3, 4^. The T1R1/T1R3 heterodimer functions as an umami taste receptor in mammals and detects l-amino acids and 5’-ribonucleotides ^5–7^. The mammalian T1R2/T1R3 heterodimer acts as a sweet sensor ^6, 8^. Likewise, homologs of *TAS1R* family genes have been identified in teleost fishes ^9^, and each of the heterodimers T1R1/T1R3 and T1R2/T1R3 can sense several amino acids in teleosts ^10^.

A previous phylogenetic analysis revealed that all mammalian and teleost *TAS1R*s can be grouped into the *TAS1R1*, *TAS1R2*, and *TAS1R3* clades ^11^, suggesting that their common ancestor had only three T1R members derived from gene duplications that have been retained in present-day species. Lineage-specific duplications and losses of *TAS1R* genes have occurred within each of these three clades, as exemplified by multiple *TAS1R2* genes in zebrafish and fugu and loss of *TAS1R2* in birds ^12^. A few genomic studies of vertebrates such as reptiles and non-teleost fishes have suggested the existence of taxonomically unplaced *TAS1R*s that may not be included in the aforementioned three clades ^13–15^. However, the lack of comprehensive characterization and systematic classification has limited our understanding of the evolutionary history of *TAS1R* genes, the functional diversity of T1Rs, and the molecular basis of taste sense in vertebrates.

Here, we present an evolutionary analysis of diverse *TAS1R*s in jawed vertebrates, with the first-ever exhaustive taxon sampling encompassing all major ‘fish’ lineages. In addition to clades *TAS1R1*, *TAS1R2*, and *TAS1R3*, we identified five novel *TAS1R* clades. The results suggest that the vertebrate ancestor possessed more T1Rs than most modern vertebrates, challenging the paradigm that only three T1R family members have been retained during evolution. Functional analyses suggest that the novel T1Rs have shaped the diversity of taste sense. We propose that the T1R family has undergone an ancient birth-and-death evolution that accelerated their functional differentiation, which may have led to the diversification of feeding habitats among vertebrates.

## Results

### Identification of novel *TAS1R* family members

We identified homologs of *TAS1R* genes that are included in public genome/transcriptome databases for diverse taxa of jawed vertebrates (Table S1). A phylogenetic analysis revealed the existence of many *TAS1R*s that had not been categorized into any of the three clades *TAS1R1*, *TAS1R2*, or *TAS1R3*; these *TAS1R*s were found in lizards, axolotl, lungfishes, coelacanth, bichir, and cartilaginous fishes (Fig. 1a, Figs. S1 and S2). These novel *TAS1R*s could be classified into five new clades. One clade, the sister clade of *TAS1R3*, was named *TAS1R4*. Clade *TAS1R4* contains genes from all the aforementioned species but not mammals, birds, crocodilians, turtles, or teleost fishes (Fig. 1b). Another novel *TAS1R*, named *TAS1R5*, was present in axolotl, lungfishes, and coelacanth and was determined to be a sister clade of the clade comprising *TAS1R1* and *TAS1R2* (Fig. 1).

**Fig. 1.**
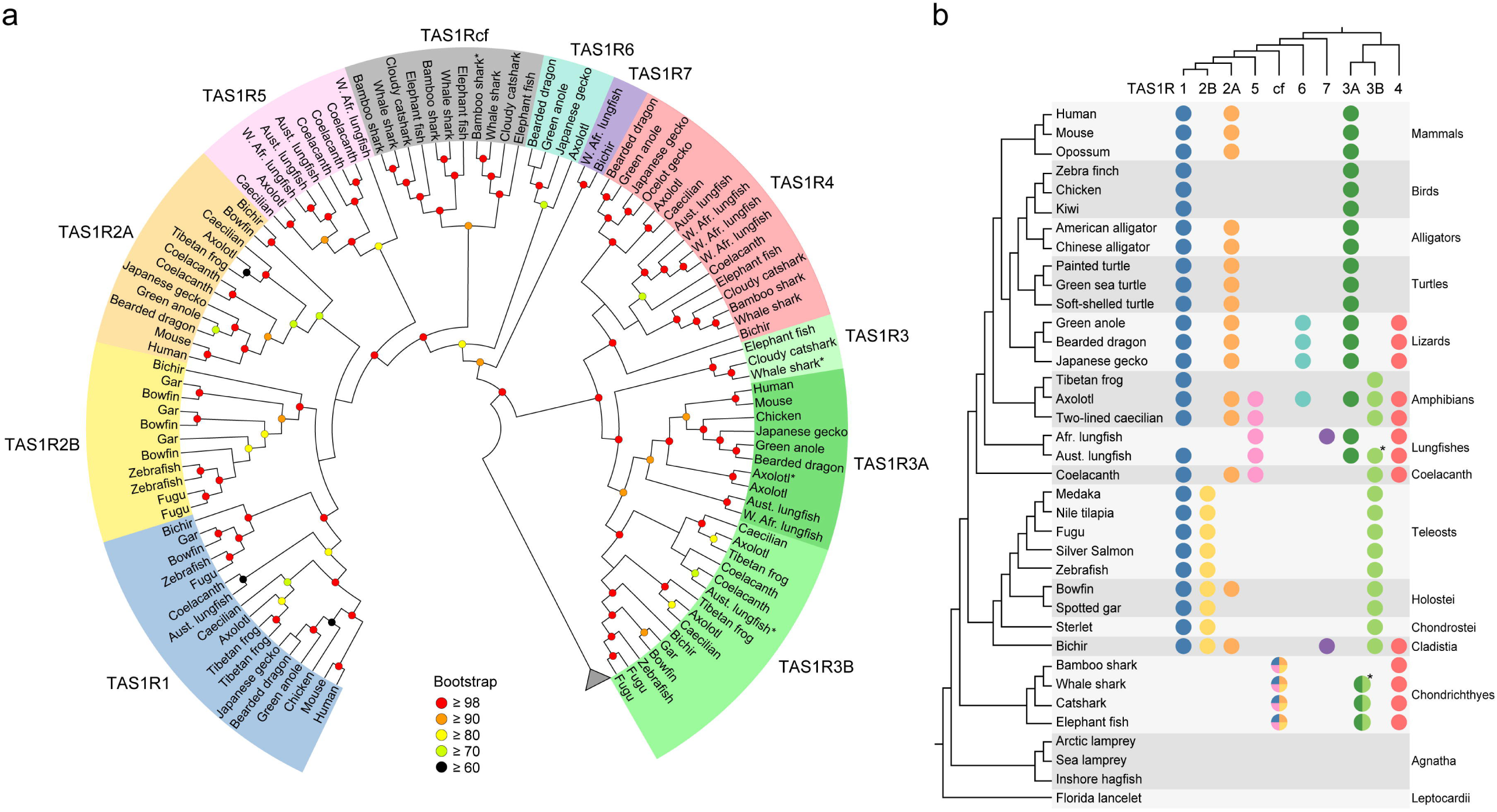
Phylogenetic tree and the revised classification of *TAS1R* members. **a,** Maximum-likelihood tree for amino acid sequences inferred from *TAS1R*s for 21 jawed vertebrates constructed with the JTT-CAT model in RAxML. Colored circles in each node represent bootstrap probabilities calculated with 1,000 replications, whereas nodes with low support (bootstrap probability < 60) have no circles. Species classification is represented with colored highlighting at the tips of the tree. GPRC6A was used as an outgroup (not shown). **b,** Distribution of *TAS1R* members among 37 chordates. The color of circles corresponds to the colored highlighting in panel ‘a’ and indicates the presence of *TAS1R* members in the genome assemblies of the various chordates. Phylogenetic relationships among species and among *TAS1Rs* are shown on the left and top, respectively. *TAS1Rcf* of cartilaginous fishes is the ortholog of the *TAS1R1*/*2A*/*2B*/*5* clade and is shown as a circle with assorted colors. Similarly, *TAS1R3* of cartilaginous fishes is shown with two shades of green that represent *TAS1R3A* and *TAS1R3B*. Circles with asterisks denote putative pseudogenes.

The sister clade to *TAS1R1* + *TAS1R2* + *TAS1R5* was identified exclusively in cartilaginous fishes (denoted *cf* in clade names) and named *TAS1Rcf*. *TAS1Rcf* could be further divided into three subclades, namely *TAS1Rcf-1*, *TAS1Rcf-2*, and *TAS1Rcf-3*, all of which were found to be present in elephant fish (also called elephant shark), belonging to the taxon Holocephali of cartilaginous fishes (Fig. S1 and S2). Therefore, the three *TAS1Rcf* subclades are likely to have emerged in the common ancestor of extant cartilaginous fishes. A thorough search of the genomes and transcriptomes of the four cartilaginous fish species identified only *TAS1R3*, *TAS1R4*, and *TAS1Rcf* but no orthologs of *TAS1R1*, *TAS1R2*, or *TAS1R5* (Fig. 1b), suggesting that *TAS1Rcf* is orthologous to the clade comprising *TAS1R1* + *TAS1R2* + *TAS1R5*.

Another novel *TAS1R* clade, *TAS1R6*, was found exclusively in axolotl and lizards. Further, the other clade, *TAS1R7*, was identified only in bichir and lungfishes, and a sister-relationship of the two genes was robustly supported (Fig. 1, Figs. S1 and S2), suggesting that *TAS1R7* emerged in their common ancestor. Indeed, the likelihood of an alternative relationship, i.e., in which *TAS1R6* and *TAS1R7* form an exclusive cluster and represent a species tree, was statistically rejected based on the approximately unbiased test (*p* < 10^−4^; Fig. S3), suggesting that *TAS1R6* and *TAS1R7* are distinct groups. Among the species included in the study, no *TAS1R* homologs were found in the two jawless vertebrates (lamprey and hagfish) or invertebrates (e.g., lancelet) (Fig. 1b).

### Each of *TAS1R3* and *TAS1R2* consists of two paralogous clades

Another remarkable finding of the phylogenetic analysis was that *TAS1R3* of bony vertebrates could be divided into two clades with high node support, which we named *TAS1R3A* and *TAS1R3B* (Fig. 1, Figs. S1 and S2). *TAS1R3A* was found to be present in tetrapods and lungfishes but not in other vertebrates, whereas *TAS1R3B* was identified only in amphibians, lungfishes, coelacanth, and ray-finned fishes. This distribution suggested that an ancestral *TAS1R3* gene was duplicated in the common ancestor of bony vertebrates, with subsequent independent loss of *TAS1R3A* in certain lineages such as coelacanth and ray-finned fishes, whereas *TAS1R3B* was lost in Amniota (mammals, birds and reptiles). Therefore, the *TAS1R3* genes in mammals and teleost fishes are paralogous to each other. Axolotl and Australian lungfish retained both *TAS1R3A* and *TAS1R3B* although the lungfish *TAS1R3B* has been pseudogenized. Furthermore, the amphibians possess two groups of *TAS1R3B* (named *TAS1R3B1* and *TAS1R3B2*; Figs. S1 and S2), suggesting that *TAS1R3B* was duplicated again at the latest before the common ancestor of amphibians.

Also, *TAS1R2* did not form a single clade in the tree (Fig. 1). The *TAS1R2* genes in ray-finned fishes formed a clade with *TAS1R1*, and the other *TAS1R2* group from tetrapods, lungfish, coelacanth, bowfin, and bichir was a sister of them. The paraphyletic relationship of the two *TAS1R2* groups is concordant with previous reports ^13^. Hereafter, we refer to the major vertebrate group as *TAS1R2A* and the ray-finned fish group as *TAS1R2B* (Fig. 1). Notably, we found that the anciently diverged ray-finned fishes such as bowfin and bichir retained both *TAS1R2A* and *TAS1R2B* as well as *TAS1R1*. We assessed the likelihood of other phylogenetic relationships in which *TAS1R2*s have a single origin, and the hypotheses were significantly rejected (*p* < 10^−6^, approximately unbiased test; Fig. S3). These results suggested that the *TAS1R2* genes in mammals and teleost fishes are probably paralogs. Thus, the *TAS1R* phylogenetic tree comprised a total of 10 *TAS1R* clades: *TAS1R1*, *TAS1R2A*, *TAS1R2B*, *TAS1R3A*, *TAS1R3B*, *TAS1R4*, *TAS1R5*, *TAS1R6*, *TAS1R7*, and *TAS1Rcf*. This unexpected gene diversity challenges the conventional conceptions about the evolution of the genetic basis for umami and sweet receptors.

### Birth-and-death evolution of the *TAS1R* family

Some of the higher-level relationships among the *TAS1R* clades were supported with relatively high node support, as exemplified by the exclusive cluster of *TAS1R3* + *TAS1R4*, the clade of the other *TAS1R*s, the clade of *TAS1R1* + *TAS1R2B* + *TAS1R2A* + *TAS1R5*, and the sister relationship of this latter clade to *TAS1Rcf* (Fig. 1, Fig. S1, Fig. S2). Based on the phylogenetic relationships and the distribution of all *TAS1R* members (Fig. 1b), the most parsimonious evolutionary scenario could be deduced as follows (Fig. 2). The first *TAS1R* gene emerged in the ancestral lineage of jawed vertebrates during the period 615–473 million years ago (Mya). This ancestral *TAS1R* underwent multiple duplications to produce at least five *TAS1R*s: *TAS1R3* (the ancestral gene of *TAS1R3A* + *TAS1R3B*), *TAS1R4*, *TAS1R6*, *TAS1R7*, and the ancestral gene of *TAS1R1* + *TAS1R2B* + *TAS1R2A* + *TAS1R5*, the latter of which corresponds to the current *TAS1Rcf* in cartilaginous fishes. In the stem lineage of bony vertebrates (473–435 Mya), *TAS1R1*, *TAS1R2A*, *TAS1R2B*, and *TAS1R5* were generated via additional gene duplication events. Simultaneously, the ancestral *TAS1R3* diverged to *TAS1R3A* and *TAS1R3B*, resulting in a total of nine *TAS1R*s in the common ancestor of bony vertebrates (Fig. 2). After the divergence of ray-finned and lobe-finned fishes ∼435 Mya, a portion of the expanded *TAS1R*s began to be differentially lost during vertebrate evolution. For example, *TAS1R7* was lost in the tetrapod ancestor, *TAS1R3B* and *TAS1R5* were lost in the amniote ancestor, and *TAS1R4* and *TAS1R6* were lost in the mammalian ancestor (Fig. 2). Thus, gene expansion before the common ancestor of bony vertebrates as well as the subsequent loss of a subset of genes have resulted in the rather dispersed distribution of *TAS1R*s in extant species (Fig. 1b).

**Fig. 2.**
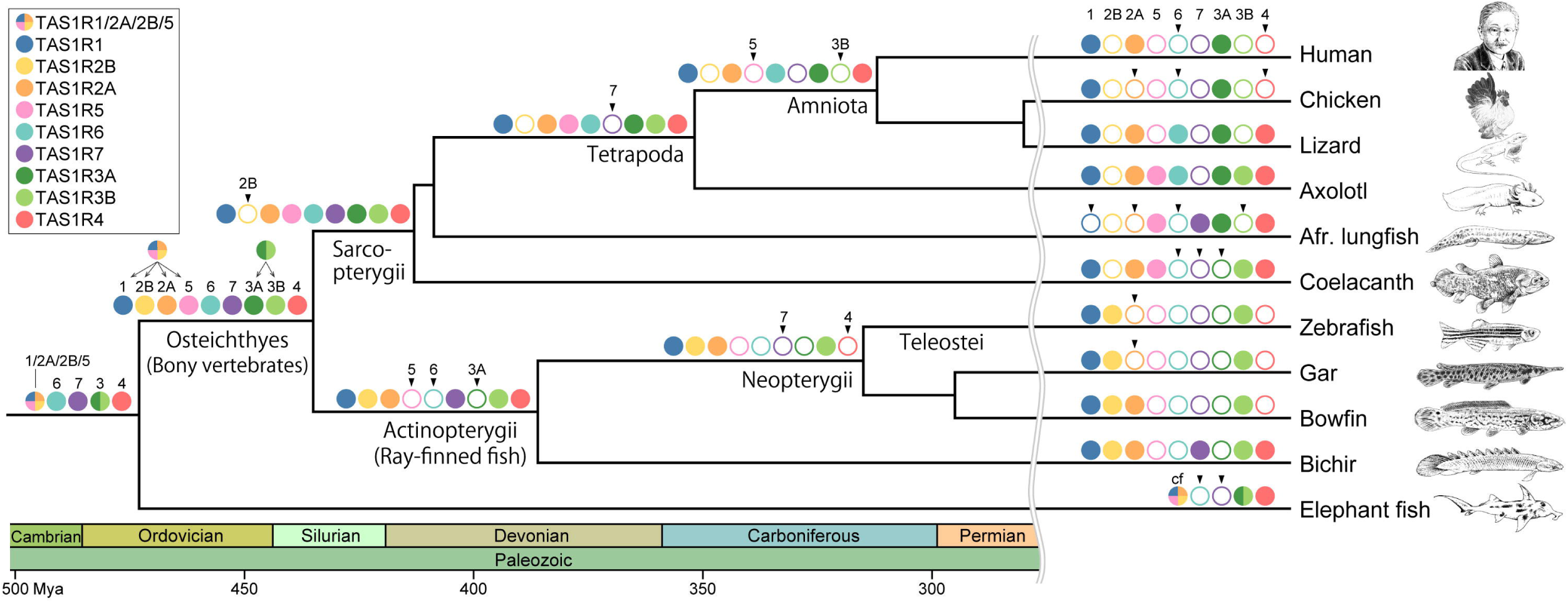
Birth-and-death history of the *TAS1R* family genes during vertebrate evolution. The color key indicates the names of the various *TAS1R* members. Filled circles on the branches indicate the presence of the *TAS1R* members, whereas open circles indicate their absence, as estimated based on the phylogenetic tree (Fig. 1a) and distribution among vertebrates (Fig. 1a). Arrowheads above open circles indicate that the *TAS1R* member was lost at the branch. Geological periods and ages (millions of years ago, Mya) are shown at the bottom. Taxon names are shown below branches. Species-specific gene duplication events for each *TAS1R* were ignored. Illustrations of the species, such as Kikunae Ikeda as a representative of humans, are shown on the right.

### *TAS1R* gene cluster retrieved by scanning understudied vertebrate genomes

The simplest model for gene amplification is a tandem duplication that produces multiple genes located side-by-side ^16, 17^. However, *TAS1R1*, *TAS1R2*, and *TAS1R3* are located far from each other in both mammalian and teleost genomes. In human chromosome 1, for example, *TAS1R1* is 12 Mb distant from *TAS1R2A* and 5 Mb distant from *TAS1R3*, with many intervening genes in each case. In zebrafish, each of *TAS1R1* and *TAS1R3B* is located on different chromosomes from the two copies of *TAS1R2*, prompting us to hypothesize that *TAS1R* members may have undergone expansion by tandem duplications in the ancestral genome, followed by subsequent translocation to distant regions during evolution. To address this possibility, the synteny of *TAS1R3* and *TAS1R4* was investigated among vertebrates, particularly those having the novel *TAS1R*s (Fig. 3, Fig. S4). Indeed, the novel *TAS1R*s were found to be located side-by-side in anole lizard, axolotl, lungfish, coelacanth, and elephant fish (Fig. 3a). Even *TAS1R2A* and *TAS1R3B* are located next to each other in axolotl and bichir. This result suggested that a *TAS1R* gene cluster had formed in the common ancestor of jawed vertebrates.

**Fig. 3.**
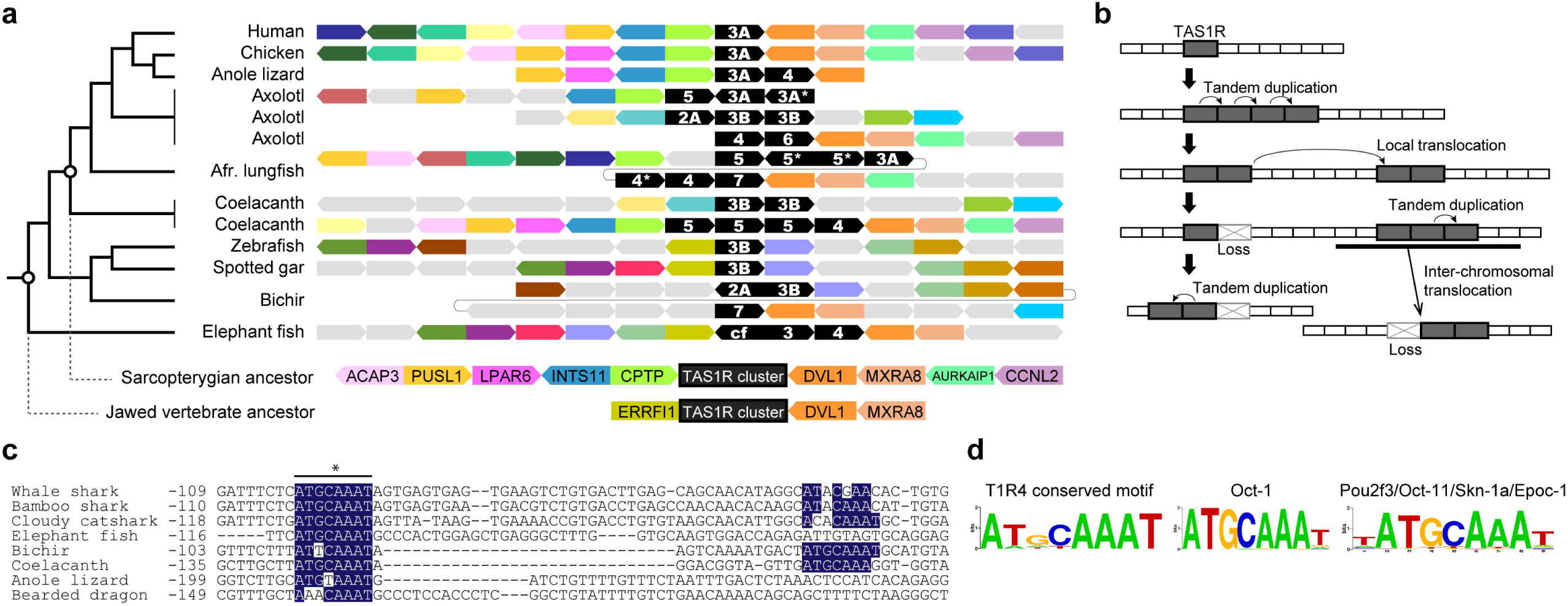
Synteny around *TAS1R*s and conserved Oct-like motifs in the *TAS1R4* upstream regions across vertebrates. **a**, Synteny around each *TAS1R* gene cluster is partly conserved across representative vertebrates. *TAS1R*s are represented by black polygons, and those with asterisks are putative pseudogenes. Colored polygons indicate genes shared among species, and gray color represents genes not shared among the species or unknown. The species tree is shown on the left. The deduced gene orders in common ancestors of Sarcopterygii and jawed vertebrates are shown at the bottom. **b**, Proposed model for the expansion of *TAS1R* genes across distant chromosomal regions during evolution. **c**, Conserved motifs located upstream of *TAS1R4*. Sequence alignment of the upstream region of the *TAS1R4* open reading frame revealed two conserved Oct-like transcription-factor binding motifs (blue shading). Numbers represent nucleotide positions from the *TAS1R4* start codon site. The asterisk indicates one of the motifs that significantly resembles the Oct-factor binding motif. **d**, Sequence logo for the conserved motif denoted with the asterisk in (c). Known binding motifs of Oct-1 (retrieved from TRANSFAC) and Oct-11/Pou2f3/Skn-1a/Epoc-1 (retrieved from JASPAR) are compared.

A comparison of neighboring genes revealed that the *TAS1R* cluster is flanked by two genes, namely *DCL1* and *MXRA8*, in the genomes of human, chicken, axolotl, lungfish, coelacanth, bichir, and elephant fish (Fig. 3a), suggesting that these two genes were adjacent to the *TAS1R* cluster in the common ancestor of jawed vertebrates. On the opposite end of the *TAS1R* cluster, the gene order of *ACAP3*–*PUSl1*–*LPAR6*–*INTS11*–*CPTP* may have been established in the sarcopterygian ancestor based on conservation among coelacanth and chicken and partly in lizard. Furthermore, the presence of other *TAS1R*-proximal genes is also conserved even across distant chromosomal regions (Fig. S4). This suggested that a chromosomal region containing both *TAS1R* and multiple neighboring genes—rather than the *TAS1R* gene alone—had translocated to a different region in each lineage. Based on the inferred ancestral gene order, the unique distribution of *TAS1R*s among present-day mammals and teleost fishes may have been a consequence of a combination of several events: 1) tandem duplication producing a *TAS1R* cluster, 2) local translocation of a subset of *TAS1R*s within a chromosome, 3) translocation of entire *TAS1R*-containing regions to different chromosomes, and 4) gene loss(es) in each lineage (Fig. 3b). Moreover, lineage-specific duplication events have occurred such as *TAS1R2B* in zebrafish and Fugu and *TAS1R2A* in coelacanth (Fig. 1a and S4) ^12, 13^. Finally, we found that some of the *TAS1R*s identified have been pseudogenized; e.g., the whale shark *TAS1R3* and the lungfish *TAS1R3B* (Fig. 1). These observations also support the evolutionary model of the *TAS1R* family presented in Fig. 3b.

### Conservation of a possible binding site for the transcription factor Oct in *TAS1R4*

Because *TAS1R4* is shared among a wide variety of vertebrates in contrast to the other novel *TAS1R*s, we expected that a transcriptional regulatory mechanism might be conserved among the species. To explore existence of a possible regulatory element, the upstream sequences of the *TAS1R4* open reading frames were aligned, and MEME^18^ was used to search for transcription-factor binding motifs conserved among the species. The most significant hit was the binding motif for the Oct family (*p* < 10^−12^ and *p* < 10^−7^ for Oct-4 and Oct-1, respectively). At least one sequence of the known Oct-binding motif ’ATGCAAAT’ is conserved among cartilaginous fishes, coelacanth, bichir, and lizards in the region upstream of *TAS1R4* (Fig. 3c, 3d). Although little is known about the transcriptional regulatory network in taste receptor cells (TRCs), one known transcription factor responsible for TRC differentiation is Skn-1a, an Oct factor also known as Oct-11, Epoc-1, or Pou2f3 ^19^. In mammals, Skn-1a is exclusively expressed in umami, sweet, and bitter TRCs, and loss of Skn-1a results in the complete absence of these taste receptor cells ^19, 20^. This finding suggested that *TAS1R4* expression is governed by a conserved regulatory mechanism involving an Oct transcription factor, possibly Skn-1a. Although Oct binding sites were not observed in the other novel *TAS1R*s, these findings may help to elucidate the molecular mechanisms underlying the conserved and/or lineage-specific expression of a variety of *TAS1R*s in TRCs.

### T1R diversity enhances the range of taste sensation

To examine which T1R receptors can form heterodimers and which ligands they respond to, we performed a cell-based functional analysis for the T1Rs of bichir, which possesses two newly discovered T1R groups (T1R4 and T1R7) and four known T1R groups (T1R1, T1R2A, T1R2B, and T1R3B). It has been proposed that T1R1 and T1R2 are responsible for ligand recognition ^21^, whereas T1R3 plays a subsidiary role, such as intersubunit conformational coupling, G-protein coupling, or membrane trafficking of T1R heterodimers ^22^. Because *TAS1R4* was found to be present in all vertebrates that harbor the other novel *TAS1R*s (Fig. 1b), T1R4 could be assumed to form a heterodimer with another T1R. We combined either T1R3B or T1R4 with another T1R (T1R1, T1R2A, T1R2B, or T1R7) in the functional analysis (Fig. 4a). Among these receptor pairs, strong responses to amino acids were detected for T1R1/T1R3B, T1R2B/T1R3B, and T1R4/T1R7 (Fig. 4b and Fig. S5). For bichir T1R2A, its combination with T1R3B or T1R4 did not yield a response to any of the tastants examined (Fig. S5a). Responses were not observed when T1R4 or T1R7 alone was used (Fig. S5a), suggesting that these newly discovered T1Rs function as an obligate heterodimers in bichir.

**Fig. 4.**
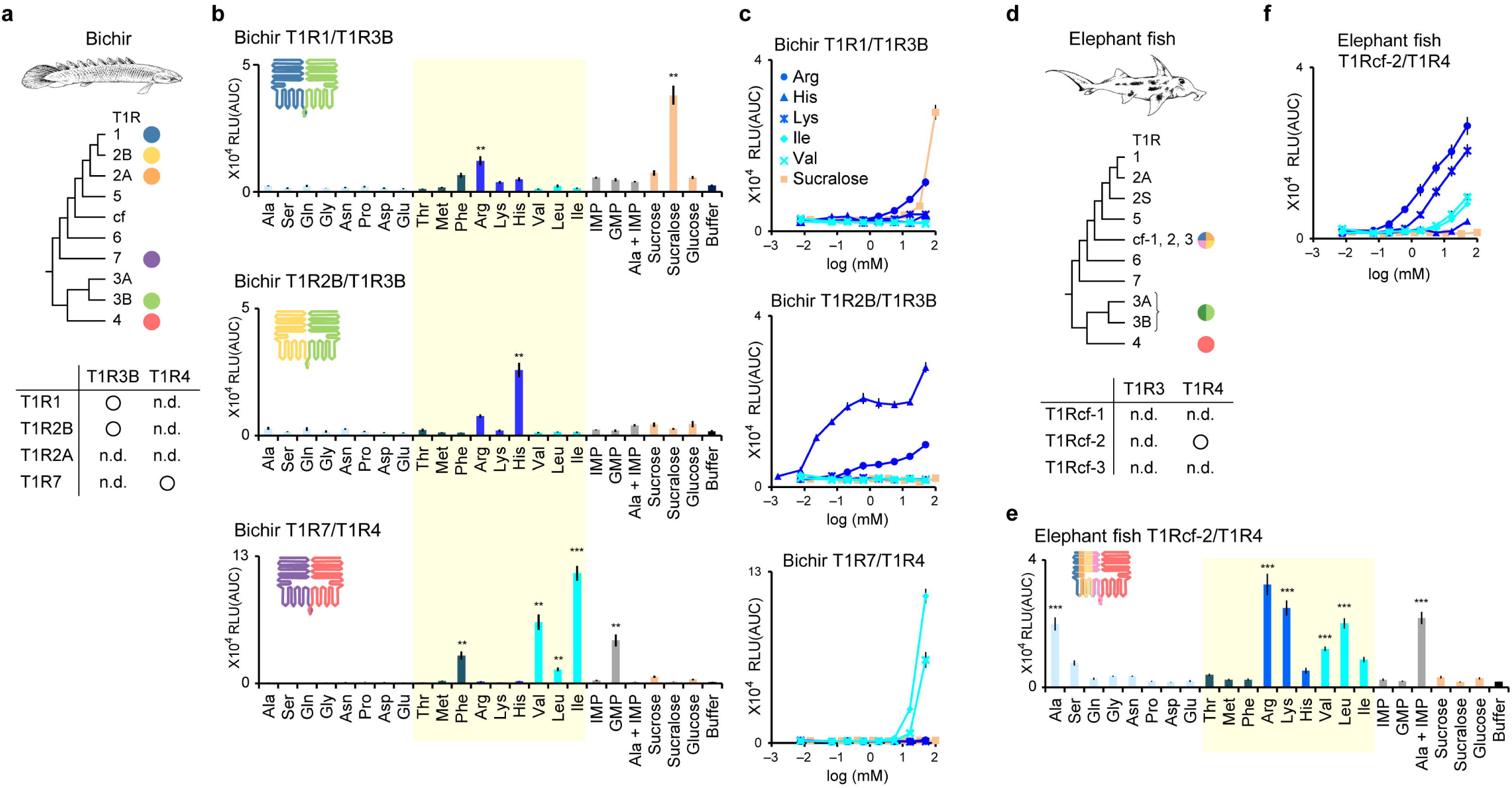
Functional analysis of T1Rs from bichir and elephant fish. **a**, T1R repertoire in bichir and their combinations used for the functional analysis. n.d.; not detected for any ligands tested. **b**, Responses of three combinations of T1R1/T1R3B (top), T1R2B/T1R3B (middle), and T1R7/T1R4 (bottom) to each of 17 amino acids (50 mM), nucleic acids (10 mM), sugars and sucralose (100 mM). Values represent the mean ± standard error of six independent experiments performed with duplicate samples. **: >10,000 relative light units with a false discovery rate (q) of <0.01; ***: >10,000 relative light units with q < 0.001. Amino acids that are essential in fishes are highlighted with yellow. **c**, Dose-response curves for T1R1/T1R3B (top), T1R2/T1R3B (middle), and T1R4/T1R7 (bottom) to three basic amino acids (Arg, His, and Lys; blue), two branched-chain amino acids (Ile and Val; light blue), and an artificial sweetener (sucralose; orange). Values represent the mean ± standard error of six independent experiments performed with duplicate samples. **d**–**f**, Same as a–c for elephant fish and the functional analysis of T1Rcf-2/T1R4.

The bichir T1R7/T1R4 responded strongly to branched-chain amino acids (BCAA; Ile, Val, and Leu) and Phe, whereas T1R1/T1R3B and T1R2B/T1R3B responded strongly to basic amino acids (Arg and His) (Fig. 4b and 4c). Fishes have 12 nutritionally essential amino acids (Cys, His, Ile, Leu, Lys, Met, Phe, Arg, Thr, Trp, Tyr, and Val) ^23^, 9 of which are included in the 17 amino acids that were tested in the T1R functional analysis. Notably, all six amino acids to which the bichir T1Rs responded are essential amino acids (*p* < 0.05; one-sided Fisher’s exact test), suggesting that the bichir T1Rs may sense essential amino acids in foods by taking advantage of the ability to perceive BCAA via the T1R4-related receptor.

Bichir T1R1/T1R3B also responded to sucralose, a structural analog of sucrose. Although only T1R2A/T1R3A is responsible for sugar perception in mammals and reptiles ^24^, we previously demonstrated that T1R1/T1R3A of birds has gained the ability to detect sugars ^25, 26^. Also, T1R2B/T1R3B of two teleost fishes, namely carp ^27^ and gilthead seabream ^28^, can detect sugars at high concentrations (100–200 mM). In addition, we found that bichir T1R7/T1R4 could respond to GMP, although a previous study reported that neither T1R1/T1R3B nor T1R2B/T1R3B of medaka fish nor T1R2B/T1R3B of zebrafish could be activated by 5’-ribonucleotides ^10^. Therefore, the origin and evolution of sugar and nucleotide taste perception may need to be reconsidered based on future genetic and functional analyses of T1Rs.

We also performed a functional analysis of elephant fish T1Rs. Three genes of the T1Rcf clade, namely T1Rcf-1, T1Rcf-2, and T1Rcf-3, were tested in combination with T1R3 and T1R4, and only the response of the T1Rcf-2/T1R4 pair could be detected (Fig. 4d–f, Fig. S5b). This combination responded to a relatively broad range of amino acids, including both BCAA (Val, Leu) and basic amino acids (Arg, Lys). The T1Rs of mammals and teleosts have little or no response to BCAA but can respond to basic amino acids ^5, 10, 29^. The observed strong response of bichir T1R7/T1R4 and elephant fish T1Rcf-2/T1R4 to BCAA may reflect functional characteristics of the novel T1Rs involving T1R4 and possibly that of ancient T1Rs in the vertebrate ancestor.

### Expression of the novel T1Rs in taste receptor cells

To investigate whether the novel T1Rs are indeed expressed in TRCs, we performed *in situ* hybridization with sections of the lips and gill rakers of bichir (Fig. 5a). T1R1, T1R2B, T1R3B, T1R4, and T1R7 were expressed in subsets of TRCs. Genes encoding downstream signal transduction molecules, such as TRPM5, Gαia1, and Gα14, were also highly expressed in subsets of TRCs in the lips and gill rakers. The signal frequencies for TRPM5, Gαia1, and Gα14 were higher than those for T1Rs.

**Fig. 5.**
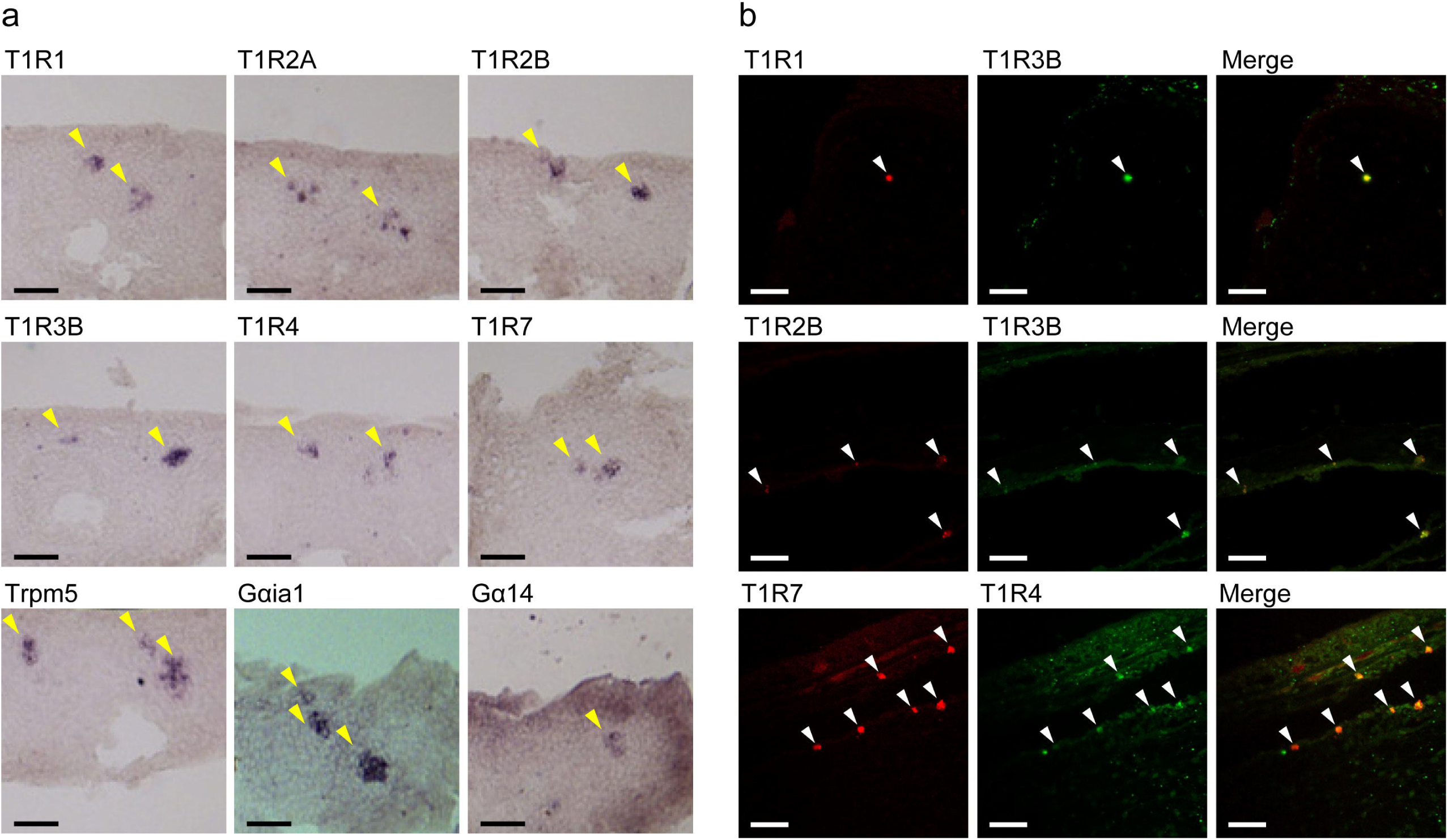
*In situ* hybridization of T1Rs in taste receptor cells of bichir. **a**, Expression of six T1Rs and three marker genes in sagittal sections. Yellow arrowheads indicate taste receptor cells expressing the various genes. Scale bar: 50 μm. **b**, Double-label fluorescence *in situ* hybridization for the combinations of T1R1/T1R3B (top), T1R2B/T1R3B (middle), and T1R7/T1R4 (bottom) in the sections. White arrowheads indicate coexpressing cells. Scale bar: 50 μm.

To examine the localization of T1Rs in TRCs, we next performed double-label fluorescence *in situ* hybridization. This analysis confirmed the overlap of the signal for T1R1 with that of T1R3B, T1R2B with T1R3B, and T1R7 with T1R4 (Fig. 5b). These results suggested that T1R1/T1R3B, T1R2B/T1R3B, and T1R7/T1R4 function as heterodimers, in accordance with the results of our functional assays.

## Discussion

The complex history of the T1R/TAS1R family includes ancient gene expansions followed by independent lineage-specific losses, which contrasts with conventional wisdom that essentially only three members were retained during evolution^11, 30^. The evolution of certain other chemoreceptors, such as the T2R (or TAS2R) bitter-taste receptor family and olfactory receptors, followed a birth-and-death process^31^. In this mode of evolution, tens or hundreds of the receptor family/superfamily genes have undergone lineage-specific extensive duplication followed by frequent gene loss via natural selection ^30^. Our results suggest that a similar process—although less extensive than what occurred for other chemoreceptors—contributed to the phylogenetic and functional expansion of the T1R family early during evolution. *TAS1R*s were not subjected to extensive birth-and-death evolution possibly because T1R ligands are limited to amino acids, sugars, and nucleotides in contrast to T2Rs and olfactory receptors that respond to a wider range of ligands/stimulants. It is also possible that the ancient expansion might have contributed to an alternate use of T1Rs in other tissues because certain G protein–coupled receptors (including T1Rs) are expressed in the gut of mammals and fishes ^32, 33^ although their functions remain unresolved.

The functional combinations of the bichir T1R7/T1R4 and the elephant fish T1Rcf-2/T1R4 suggest that T1R4 may have a similar role to T1R3 by forming a functional heterodimer with another novel T1R such as T1R5, T1R6, T1R7, or T1Rcf. This model is also supported by the fact that species with either *TAS1R5*, *TAS1R6*, *TAS1R7*, or *TAS1Rcf* also have *TAS1R4* (Fig. 1b) and that *TAS1R4* is phylogenetically the sister group of *TAS1R3* (Fig. 1a). Therefore, the common ancestor of bony vertebrates with at least nine T1Rs likely had two types of heterodimeric T1R receptors, namely T1R3- and T1R4-dependent receptors. This relatively wide variety of possible T1R combinations involving two duplicated genes of T1R2 (A and B) and T1R3 (A and B) might have contributed to the diversification of taste sensation.

Our findings provoke new questions, one of which is why many *TAS1R* genes—particularly the T1R4-related receptors—have been frequently lost and many species have come to rely predominantly on T1R3-dependent receptors (Fig. 2). One possible reason is that the loss of one or more T1Rs might have been triggered by dietary changes that occurred in the ancestral lineages. This is plausible because previous studies reported losses of *TAS1R*s and *TAS2R*s in many land vertebrates, presumably owing to specific dietary shifts ^34–36^. Also, the behavior of swallowing foods whole, i.e., without mastication, might have reduced the necessity for taste sense, which may have led to T1R loss, as previously discussed with respect to mammals ^35, 37^ and reptiles ^38^. Alternatively, it is possible that T1R3-dependent receptors have acquired greater functional flexibility and/or evolvability than other T1Rs; i.e., various tastants might have been detected by evolutionary tuning of the sequences and structures of the T1R3-dependent receptors rather than additional gene duplication. Such cases are indeed known for land vertebrates such as primates ^7^ and birds ^25, 26^. To address these issues, it will be essential to carry out functional analyses of the newly discovered T1Rs in addition to the known T1R1/T1R3 and T1R2/T1R3 for a broad range of vertebrates, as our current results demonstrate. For example, the response to BCAA is a previously unreported characteristic shared between the bichir T1R7/T1R4 and elephant fish T1Rcf-2/T1R4 (Fig. 4). These results will provide the first insight into the sensory characteristics of an ancestor of vertebrates. The bichir T1Rs also responded to other essential amino acids, a sucrose analog, and a nucleotide. Future analysis will resolve whether the functions indeed reflect the characteristics of the ancestral species.

Thus, by demonstrating the unexpected diversity and unique evolutionary process of the T1R family, our results set the stage for understanding the evolutionary-scale changes in taste sense in vertebrates. Our understanding of taste sense will be further enhanced by clarifying T1R repertoires in each species, their tissue-specific expression, transcriptional regulatory mechanisms, and protein structures. Revealing the functional and structural diversity of the novel T1Rs will help us elucidate the molecular mechanism(s) by which human T1Rs recognize palatable tastes.

## Materials and Methods

### Identification of *TAS1R* genes from the genome and RNA-seq data in vertebrates

We used genome and transcriptome data as well as related raw sequence reads for a broad range of vertebrates (Table S1). A tblastn search was carried out using amino acid sequences of the validated *TAS1R*s in human, chicken, and zebrafish as queries. In the blast hit scaffolds, exon regions were predicted using AUGUSTUS ver. 3.2.3 ^39^, followed by an evaluation of the exon-intron boundaries by aligning the genome sequences with the query *TAS1R* sequences and by the GT/AG rule. Because frequent base errors were observed in the genome assembly for axolotl, sequence correction was needed for our *TAS1R* identification. We retrieved the raw reads of the public genome data and RNA sequencing data corresponding to the *TAS1R* exons using bowtie2 ^40^ and blastn and used that data to correct the *TAS1R* sequences by checking the alignment. The *TAS1R* amino acid sequences identified for axolotl, coelacanth, and bichir were used as queries for an additional tblastn search of other vertebrates.

### Phylogenetic analysis

Amino acid sequences of T1Rs were aligned using MAFFT ver. 7.427 with the ginsi option ^41^, followed by manual adjustment. Hypervariable and unalignable regions were removed using prequal ^42^, and a maximum likelihood tree was constructed using RAxML ver. 8.2.12 with the JTT-CAT model with 1,000 bootstrap replicates ^43^. The G protein–coupled receptor family C group 6 member A (GPRC6A) genes, which are the closest relative of T1Rs ^44^, were used as the outgroup. Bayesian tree inference was conducted with MrBayes 3.2.6 with the JTT-F + Γ_4_ model ^45^. Two simultaneous runs were carried out with 10,000,000 generations, of which 2,500,000 were discarded as burn-in, and convergence was assessed with Tracer ^46^. Trees were visualized with iTOL ^47^. Alternative tree topologies were evaluated with the approximately unbiased test using CONSEL v0.20 ^48^.

### Synteny analysis

The synteny of genes proximal to the novel T1Rs was analyzed using annotations available in Ensembl 97 ^49^ for human (GRCh38), chicken (GRCg6a), anole lizard (AnoCar2.0), coelacanth (LatCha1), zebrafish (GRCz11), and spotted gar (LepOcu1). For bichir, gene annotation data generated by Cufflinks, which will be published elsewhere, was used for our synteny analysis. The gene annotation for axolotl was obtained from the Axolotl-omics website (AmexG_3.0.0) ^50^. NCBI annotation was referred to for the West African lungfish (PAN1.0) and elephant fish (Callorhinchus_milii-6.1.3). Novel *TAS1R*s were added if they were not accurately identified in the public annotation data.

### Survey of conserved motifs in the sequence upstream of *TAS1R4*

Sequences up to 300 bp upstream of the *TAS1R4* open reading frames were collected for whale shark, bamboo shark, cloudy catshark, elephant fish, bichir, coelacanth, axolotl, two-lined caecilian, Japanese gecko, anole lizard, and central bearded dragon. The sequences were aligned using MAFFT ^41^ and then used for MEME analysis ^18^ to search for a maximum of three conserved sequence motifs. The motifs discovered by MEME were then used for comparison with known transcription-factor binding motifs in TRANSFAC v11.3 using STAMP ^51^. The known Oct-11/Pou2f3 motif was obtained from JASPAR ^52^.

### Cloning fish *TAS1R*s

*TAS1R1*, *TAS1R2A*, *TAS1R2B*, *TAS1R3B*, *TAS1R4*, and *TAS1R7* were amplified by PCR from the genomic DNA or cDNA of bichir (*Polypterus senegalus*). *TAS1Rcf-1*, *TAS1Rcf-2*, *TAS1Rcf-3*, *TAS1R3*, and *TAS1R4* were amplified by PCR from the genomic DNA of elephant fish (*Callorhinchus milii*). PCR and Sanger sequencing for the coding sequences of their *TAS1R* genes were performed using specific primers designed based on the annotation from the whole genome assemblies. The PCR products of the exons were assembled into one full-length sequence using overlapping PCR (In-fusion cloning, Clontech) for each *TAS1R* and were then subcloned into the pEAK10 expression vector (Edge Biosystems, Gaithersburg, MD).

### Functional analysis of T1Rs

Responses of the T1Rs to various taste-associated stimulants were measured by using a heterologous expression system ^29^. Briefly, HEK293T cells were transiently co-transfected with an expression vector for an individual T1R along with rat G15i2 and mt-apoclytin-II and then exposed to taste stimuli, and luminescence intensity was measured using a FlexStation 3 microplate reader (Molecular Devices). The response in each well was calculated based on the area under the curve and expressed as relative light units. Data were collected from three independent experiments carried out in duplicate. We adapted a strict definition for the positive response as over 10,000 relative light units and statistically significant differences against control (buffer) with a false discovery rate (q) of <0.01 (one-sided *t*-test).

### *In situ* hybridization

*In situ* hybridization was performed as previously described ^9^. In brief, fresh-frozen sections (10-μm thick) of bichir jaw tissue were placed on MAS-coated glass slides (Matsunami Glass, Osaka, Japan) and fixed with 4% paraformaldehyde in phosphate-buffered saline. Prehybridization (58°C, 1 h), hybridization (58°C, for two overnights), washing (58°C, 0.2× saline-sodium citrate), and development (NBT-BCIP) were performed using digoxigenin-labeled probes. Images of stained sections were obtained using a fluorescence microscope (DM6 B, Leica, Nussloch, Germany) equipped with a cooled CCD digital camera (DFC7000 T, Leica). Double-label fluorescence *in situ* hybridization was performed using digoxigenin- and fluorescein-labeled RNA probes. Each labeled probe was sequentially detected by incubation with a peroxidase-conjugated antibody against digoxigenin and peroxidase-conjugated anti-fluorescein (Roche, Indianapolis, IN, USA) followed by incubation with TSA-Alexa Fluor 555 and TSA-Alexa Fluor 488 (Invitrogen, Carlsbad, CA, USA) using the tyramide signal amplification method. Images of stained sections were obtained using a confocal laser-scanning microscope (LSM 800; ZEISS, Oberkochen, Germany). The entire coding regions for the six T1Rs and two G protein α subunits as well as the partial coding region for Trpm5, all of which were amplified from bichir cDNA synthesized from lip tissue, were used as probes for *in situ* hybridization.

## Acknowledgments

We thank Dr. Susumu Hyodo (The University of Tokyo) for providing the *Callorhinchus milii* sample. We also thank Erina Kamiya (School of Life Science and Technology, Tokyo Institute of Technology) for technical assistance. The authors acknowledge Open Facility Center, Tokyo Institute of Technology, for sequencing assistance. Computations were partially performed on the supercomputer systems at the ROIS National Institute of Genetics and the Institute of Statistical Mathematics. This study was supported by JSPS KAKENHI Grant Number 19H03272 (to H.N.), 18K14427, 20H02941, and 23H02168 (to Y.T.), Research Project Grant(B) by Institute of Science and Technology, Meiji University (to Y.I.), and the Lotte Shigemitsu Prize (to Y.T. and Y.I.).

## Author contributions

H.N., Y.T., and Y.I. conceived and supervised the study. H.N., T.K., S.K., and M.O. analyzed the vertebrate genomes. H.N. performed the phylogenetic and synteny analyses. Y.T. performed the functional assay. K.K., A.G., K.H., S.O., and Y.I. performed *in situ* hybridization experiments. H.N., Y.T., and Y.I. wrote the original draft of the manuscript. H.N., Y.T., Y.I., S.K., and M.O. edited the manuscript.

## References

1 Trivedi, B. P. Gustatory system: the finer points of taste. Nature 486, S2–3, doi:10.1038/486S2a (2012).

2 Yarmolinsky, D. A., Zuker, C. S. & Ryba, N. J. Common sense about taste: from mammals to insects. Cell 139, 234–244, doi:10.1016/j.cell.2009.10.001 (2009).

3 Li, X. et al. Human receptors for sweet and umami taste. Proc Natl Acad Sci U S A 99, 4692–4696, doi:10.1073/pnas.072090199 (2002).

4 Hummel, T. & Welge-Lüssen, A. Taste and smell : an update. (Karger, 2006).

5 Nelson, G. et al. An amino-acid taste receptor. Nature 416, 199–202, doi:10.1038/nature726 (2002).

6 Zhao, G. Q. et al. The receptors for mammalian sweet and umami taste. Cell 115, 255–266, doi:10.1016/s0092-8674(03)00844-4 (2003).

7 Toda, Y. et al. Evolution of the primate glutamate taste sensor from a nucleotide sensor. Curr Biol 31, 4641–4649 e4645, doi:10.1016/j.cub.2021.08.002 (2021).

8 Nelson, G. et al. Mammalian sweet taste receptors. Cell 106, 381–390, doi:10.1016/s0092-8674(01)00451-2 (2001).

9 Ishimaru, Y. et al. Two families of candidate taste receptors in fishes. Mech Dev 122, 1310–1321, doi:10.1016/j.mod.2005.07.005 (2005).

10 Oike, H. et al. Characterization of ligands for fish taste receptors. J Neurosci 27, 5584–5592, doi:10.1523/JNEUROSCI.0651-07.2007 (2007).

11 Shi, P. & Zhang, J. Contrasting modes of evolution between vertebrate sweet/umami receptor genes and bitter receptor genes. Mol Biol Evol 23, 292–300, doi:10.1093/molbev/msj028 (2006).

12 Bachmanov, A. A. et al. Genetics of taste receptors. Curr Pharm Des 20, 2669–2683, doi:10.2174/13816128113199990566 (2014).

13 Picone, B. et al. Taste and odorant receptors of the coelacanth--a gene repertoire in transition. J Exp Zool B Mol Dev Evol 322, 403–414, doi:10.1002/jez.b.22531 (2014).

14 Hara, Y. et al. Madagascar ground gecko genome analysis characterizes asymmetric fates of duplicated genes. BMC Biol 16, 40, doi:10.1186/s12915-018-0509-4 (2018).

15 Sharma, K., Syed, A. S., Ferrando, S., Mazan, S. & Korsching, S. I. The Chemosensory Receptor Repertoire of a True Shark Is Dominated by a Single Olfactory Receptor Family. Genome Biol Evol 11, 398–405, doi:10.1093/gbe/evz002 (2019).

16 Ohno, S. *Evolution by Gene Duplication*. (Springer, Berlin, Heidelberg, 1970).

17 Lewis, E. B. A gene complex controlling segmentation in Drosophila. Nature 276, 565–570, doi:10.1038/276565a0 (1978).

18 Bailey, T. L., Johnson, J., Grant, C. E. & Noble, W. S. The MEME Suite. Nucleic Acids Res 43, W39–49, doi:10.1093/nar/gkv416 (2015).

19 Matsumoto, I., Ohmoto, M., Narukawa, M., Yoshihara, Y. & Abe, K. Skn-1a (Pou2f3) specifies taste receptor cell lineage. Nat Neurosci 14, 685–687, doi:10.1038/nn.2820 (2011).

20 Yamashita, J., Ohmoto, M., Yamaguchi, T., Matsumoto, I. & Hirota, J. Skn-1a/Pou2f3 functions as a master regulator to generate Trpm5-expressing chemosensory cells in mice. PLoS One 12, e0189340, doi:10.1371/journal.pone.0189340 (2017).

21 Zhang, F. et al. Molecular mechanism for the umami taste synergism. Proc Natl Acad Sci U S A 105, 20930–20934, doi:10.1073/pnas.0810174106 (2008).

22 Nuemket, N. et al. Structural basis for perception of diverse chemical substances by T1r taste receptors. Nat Commun 8, 15530, doi:10.1038/ncomms15530 (2017).

23 Hou, Y. & Wu, G. Nutritionally Essential Amino Acids. Adv Nutr 9, 849–851, doi:10.1093/advances/nmy054 (2018).

24 Liang, Q. et al. T1R2-mediated sweet sensing in a lizard. Curr Biol 32, R1302–R1303, doi:10.1016/j.cub.2022.10.061 (2022).

25 Baldwin, M. W., et al. Sensory biology. Evolution of sweet taste perception in hummingbirds by transformation of the ancestral umami receptor. Science 345, 929–933, doi:10.1126/science.1255097 (2014).

26 Toda, Y. et al. Early origin of sweet perception in the songbird radiation. Science 373, 226–231, doi:10.1126/science.abf6505 (2021).

27 Yuan, X. C. et al. Expansion of sweet taste receptor genes in grass carp (Ctenopharyngodon idellus) coincided with vegetarian adaptation. BMC Evol Biol 20, 25, doi:10.1186/s12862-020-1590-1 (2020).

28 Angotzi, A. R., Puchol, S., Cerda-Reverter, J. M. & Morais, S. Insights into the Function and Evolution of Taste 1 Receptor Gene Family in the Carnivore Fish Gilthead Seabream (Sparus aurata). Int J Mol Sci 21, doi:10.3390/ijms21207732 (2020).

29 Toda, Y. et al. Two distinct determinants of ligand specificity in T1R1/T1R3 (the umami taste receptor). J Biol Chem 288, 36863–36877, doi:10.1074/jbc.M113.494443 (2013).

30 Nei, M., Niimura, Y. & Nozawa, M. The evolution of animal chemosensory receptor gene repertoires: roles of chance and necessity. Nat Rev Genet 9, 951–963, doi:10.1038/nrg2480 (2008).

31 Nei, M. & Rooney, A. P. Concerted and birth-and-death evolution of multigene families. Annu Rev Genet 39, 121–152, doi:10.1146/annurev.genet.39.073003.112240 (2005).

32 Jang, H. J. et al. Gut-expressed gustducin and taste receptors regulate secretion of glucagon-like peptide-1. Proc Natl Acad Sci U S A 104, 15069–15074, doi:10.1073/pnas.0706890104 (2007).

33 Calo, J. et al. First evidence for the presence of amino acid sensing mechanisms in the fish gastrointestinal tract. Sci Rep 11, 4933, doi:10.1038/s41598-021-84303-9 (2021).

34 Antinucci, M. & Risso, D. A Matter of Taste: Lineage-Specific Loss of Function of Taste Receptor Genes in Vertebrates. Front Mol Biosci 4, 81, doi:10.3389/fmolb.2017.00081 (2017).

35 Jiang, P. et al. Major taste loss in carnivorous mammals. Proc Natl Acad Sci U S A 109, 4956–4961, doi:10.1073/pnas.1118360109 (2012).

36 Liu, G. et al. Differentiated adaptive evolution, episodic relaxation of selective constraints, and pseudogenization of umami and sweet taste genes TAS1Rs in catarrhine primates. Front Zool 11, 79, doi:10.1186/s12983-014-0079-4 (2014).

37 Feng, P., Zheng, J., Rossiter, S. J., Wang, D. & Zhao, H. Massive losses of taste receptor genes in toothed and baleen whales. Genome Biol Evol 6, 1254–1265, doi:10.1093/gbe/evu095 (2014).

38 Feng, P. & Liang, S. Molecular evolution of umami/sweet taste receptor genes in reptiles. PeerJ 6, e5570, doi:10.7717/peerj.5570 (2018).

39 Stanke, M. & Morgenstern, B. AUGUSTUS: a web server for gene prediction in eukaryotes that allows user-defined constraints. Nucleic Acids Res 33, W465–467, doi:10.1093/nar/gki458 (2005).

40 Langmead, B. & Salzberg, S. L. Fast gapped-read alignment with Bowtie 2. Nat Methods 9, 357–359, doi:10.1038/nmeth.1923 (2012).

41 Katoh, K. & Standley, D. M. MAFFT multiple sequence alignment software version 7: improvements in performance and usability. Mol Biol Evol 30, 772–780, doi:10.1093/molbev/mst010 (2013).

42 Whelan, S., Irisarri, I. & Burki, F. PREQUAL: detecting non-homologous characters in sets of unaligned homologous sequences. Bioinformatics 34, 3929–3930, doi:10.1093/bioinformatics/bty448 (2018).

43 Stamatakis, A. RAxML version 8: a tool for phylogenetic analysis and post-analysis of large phylogenies. Bioinformatics 30, 1312–1313, doi:10.1093/bioinformatics/btu033 (2014).

44 Kuang, D. et al. Ancestral reconstruction of the ligand-binding pocket of Family C G protein-coupled receptors. Proc Natl Acad Sci U S A 103, 14050–14055, doi:10.1073/pnas.0604717103 (2006).

45 Ronquist, F. et al. MrBayes 3.2: efficient Bayesian phylogenetic inference and model choice across a large model space. Syst Biol 61, 539–542, doi:10.1093/sysbio/sys029 (2012).

46 Rambaut, A., Drummond, A. J., Xie, D., Baele, G. & Suchard, M. A. Posterior Summarization in Bayesian Phylogenetics Using Tracer 1.7. Syst Biol 67, 901–904, doi:10.1093/sysbio/syy032 (2018).

47 Letunic, I. & Bork, P. Interactive Tree Of Life (iTOL) v5: an online tool for phylogenetic tree display and annotation. Nucleic Acids Res 49, W293–W296, doi:10.1093/nar/gkab301 (2021).

48 Shimodaira, H. & Hasegawa, M. CONSEL: for assessing the confidence of phylogenetic tree selection. Bioinformatics 17, 1246–1247, doi:10.1093/bioinformatics/17.12.1246 (2001).

49 Cunningham, F., et al. Ensembl 2022. Nucleic Acids Res 50, D988–D995, doi:10.1093/nar/gkab1049 (2022).

50 Nowoshilow, S. & Tanaka, E. M. Introducing www.axolotl-omics.org - an integrated -omics data portal for the axolotl research community. Exp Cell Res 394, 112143, doi:10.1016/j.yexcr.2020.112143 (2020).

51 Mahony, S. & Benos, P. V. STAMP: a web tool for exploring DNA-binding motif similarities. Nucleic Acids Res 35, W253–258, doi:10.1093/nar/gkm272 (2007).

52 Castro-Mondragon, J. A., et al. JASPAR 2022: the 9th release of the open-access database of transcription factor binding profiles. Nucleic Acids Res 50, D165–D173, doi:10.1093/nar/gkab1113 (2022).

