## Supplementary material for "Latent taste diversity revealed by a vertebrate-wide catalogue of T1R receptors": Fig. S1-S5, Table S1

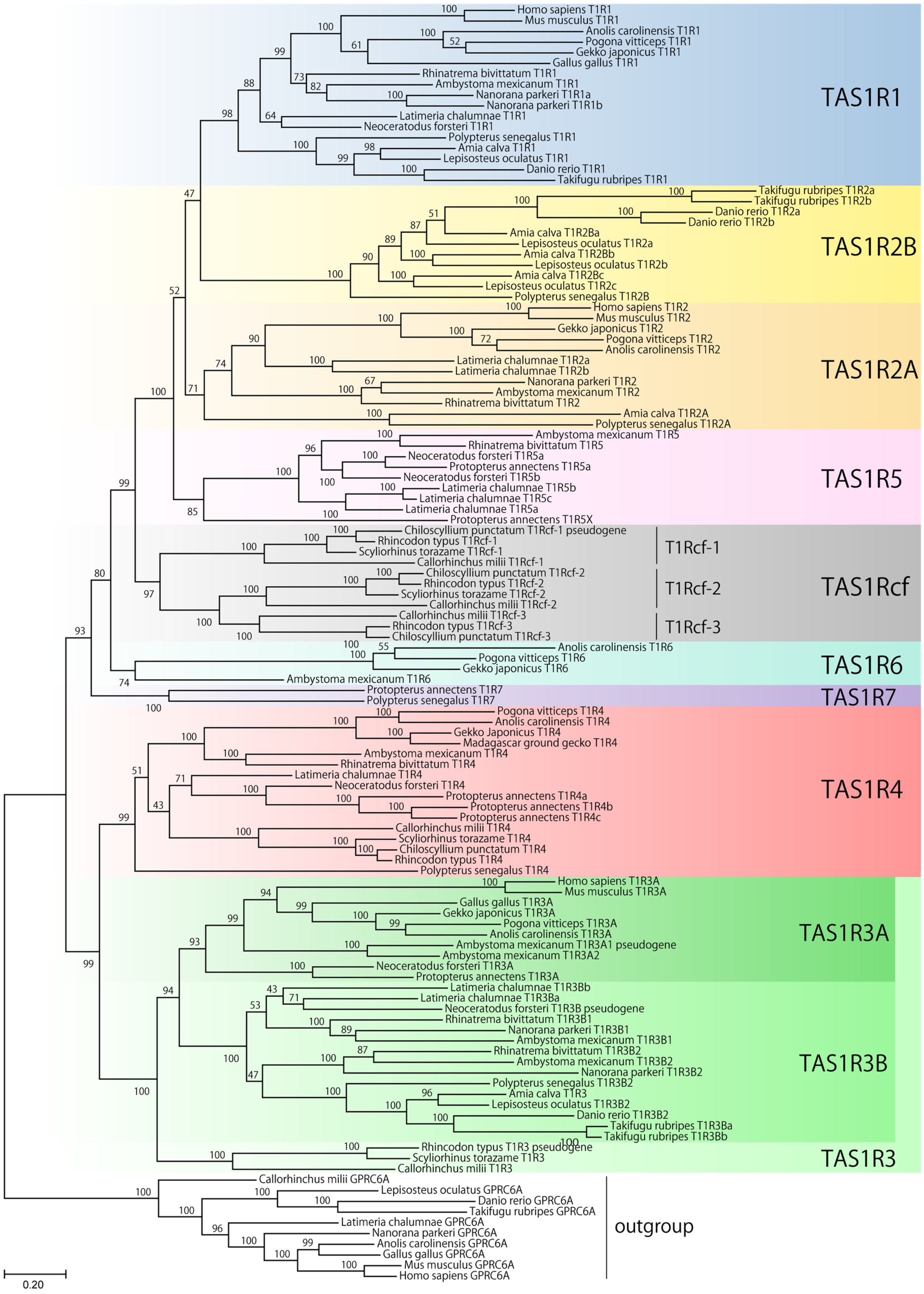


**Fig. S1. Maximum-likelihood tree of *TAS1R* members identified from 21 vertebrates.**

A maximum-likelihood tree was reconstructed from the amino acid sequences encoded by *TAS1R*s using RAxML with the JTT-CAT model. Node supports represent bootstrap probabilities calculated with 1,000 replications. *TAS1R* clade names are shown on the right.

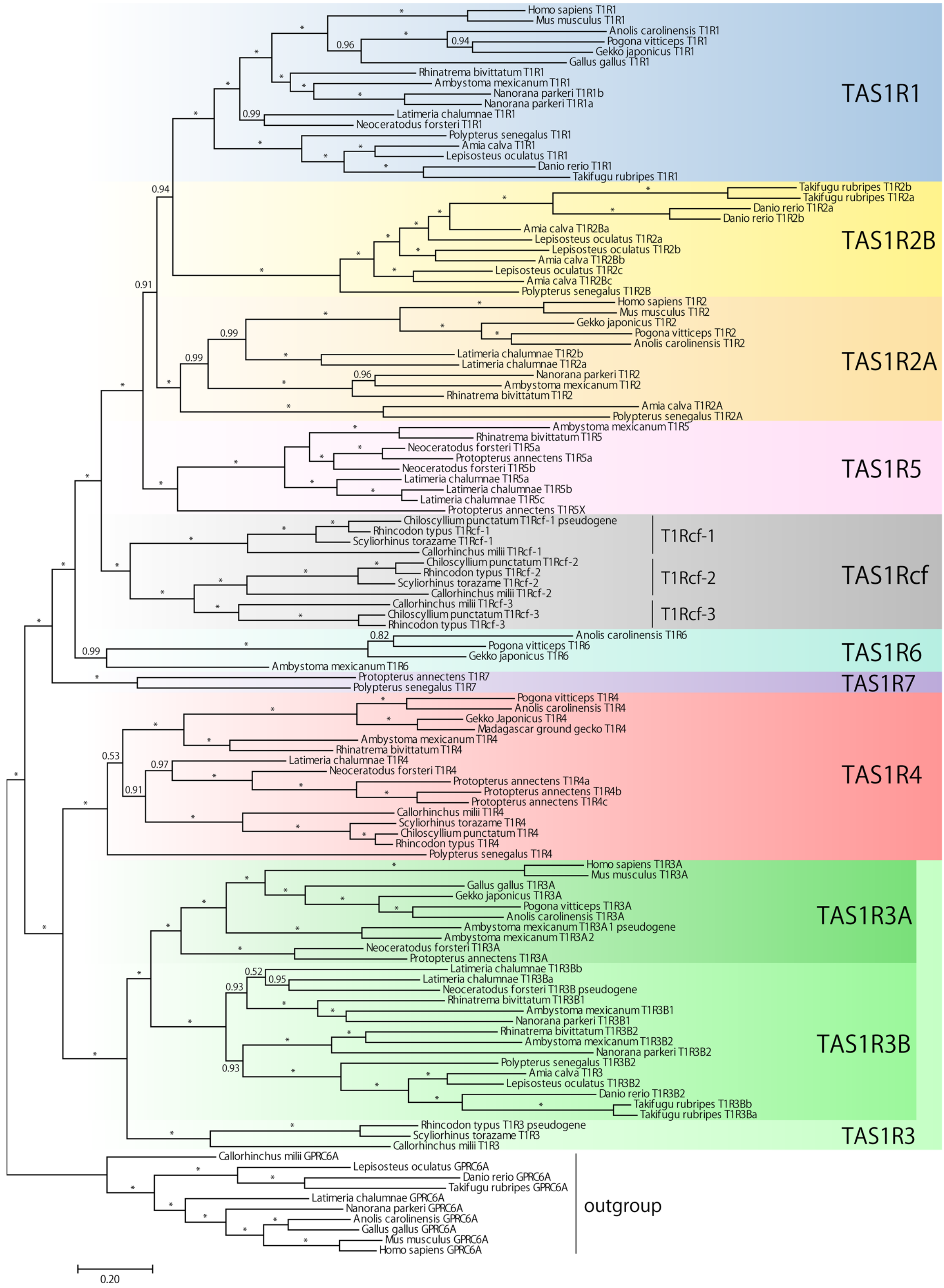


**Fig. S2. Bayesian tree of *TAS1R* members identified from 21 vertebrates.**

Bayesian tree inference was performed for the amino acid sequences encoded by *TAS1R*s using MrBayes with the JTT-F + $\Gamma_4$ model. Node supports represent Bayesian posterior probabilities, and asterisks indicate a posterior probability of 1.00. *TAS1R* clade names are shown on the right.

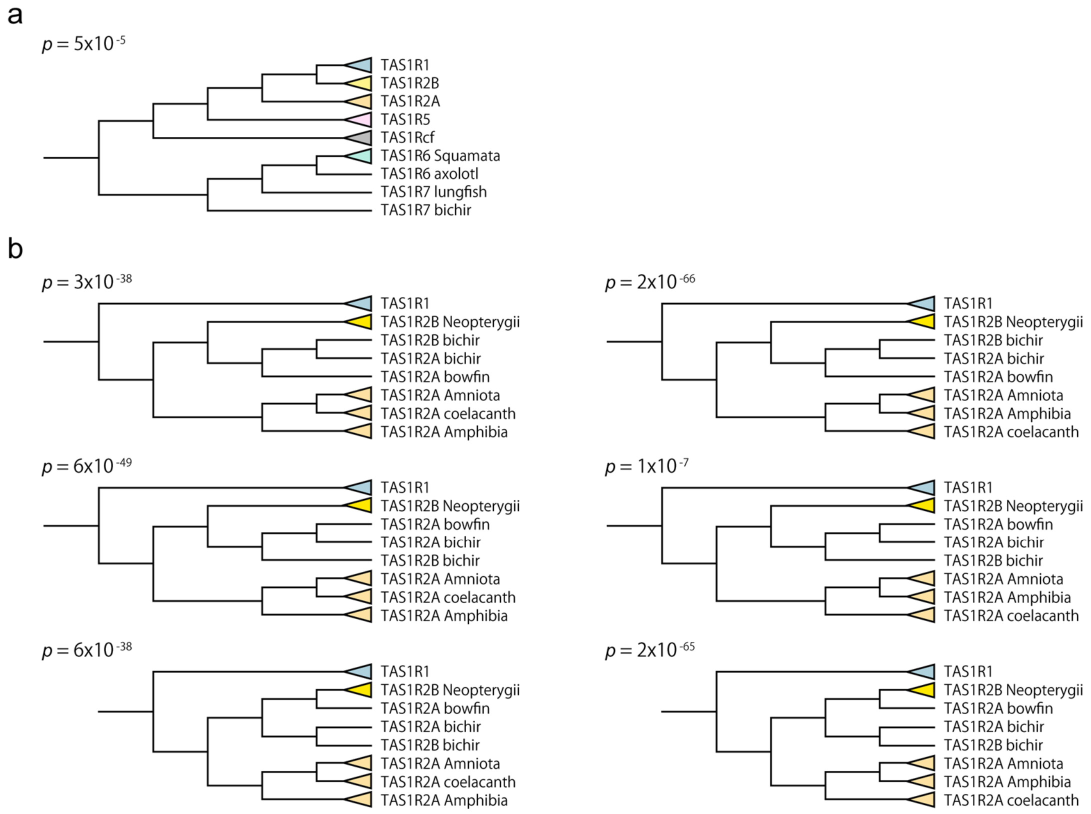


**Fig. S3. Tree topologies examined for the approximately unbiased test.**

**a**, Tree topology assuming the grouping of *TAS1R6* and *TAS1R7* and showing a species tree. **b**, Tree topologies for various relationships among *TAS1R2A* and *TAS1R2B* genes from fishes and amphibians. The *p*-values for the approximately unbiased test calculated with CONSEL are shown above each tree. Relationships within each collapsed group were fixed to be the same as in the maximum-likelihood tree (Fig. 1).

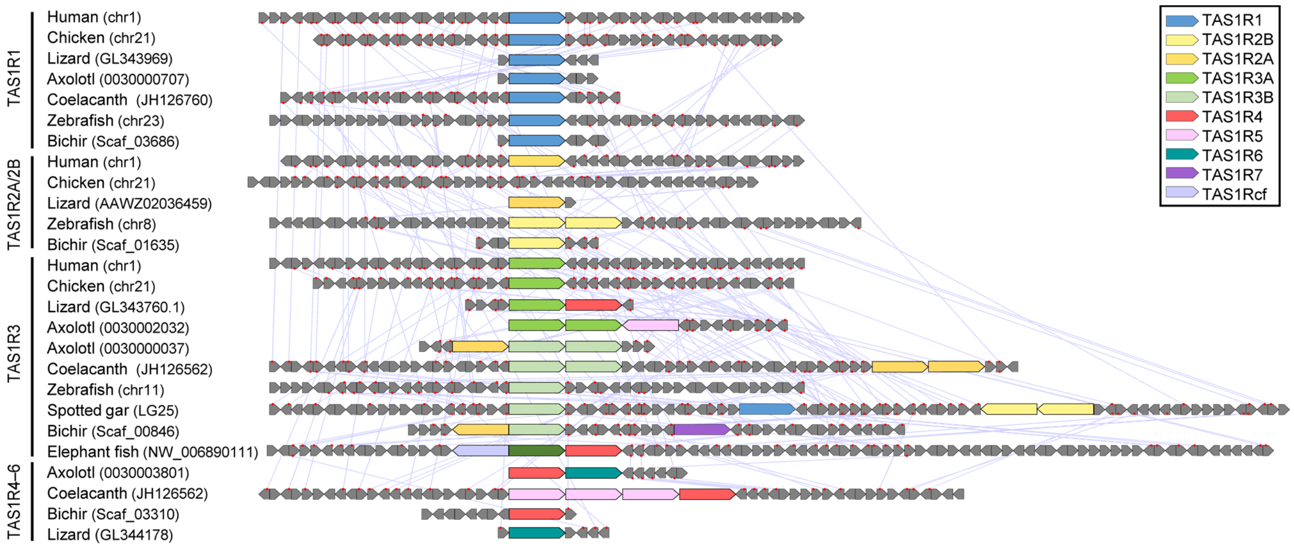


**Fig. S4. Comparison of synteny among vertebrates.**

*TAS1R*s and non-*TAS1R* genes are represented by colored and grey polygons, respectively, with each pointed end indicating the direction of transcription. Genomic regions are categorized according to the *TAS1R1*-, *TAS1R2*-, *TAS1R3*-containing regions as well as the other *TAS1R*-containing regions. Orthologous non-*TAS1R* genes between the closely represented species are connected by light-blue lines.

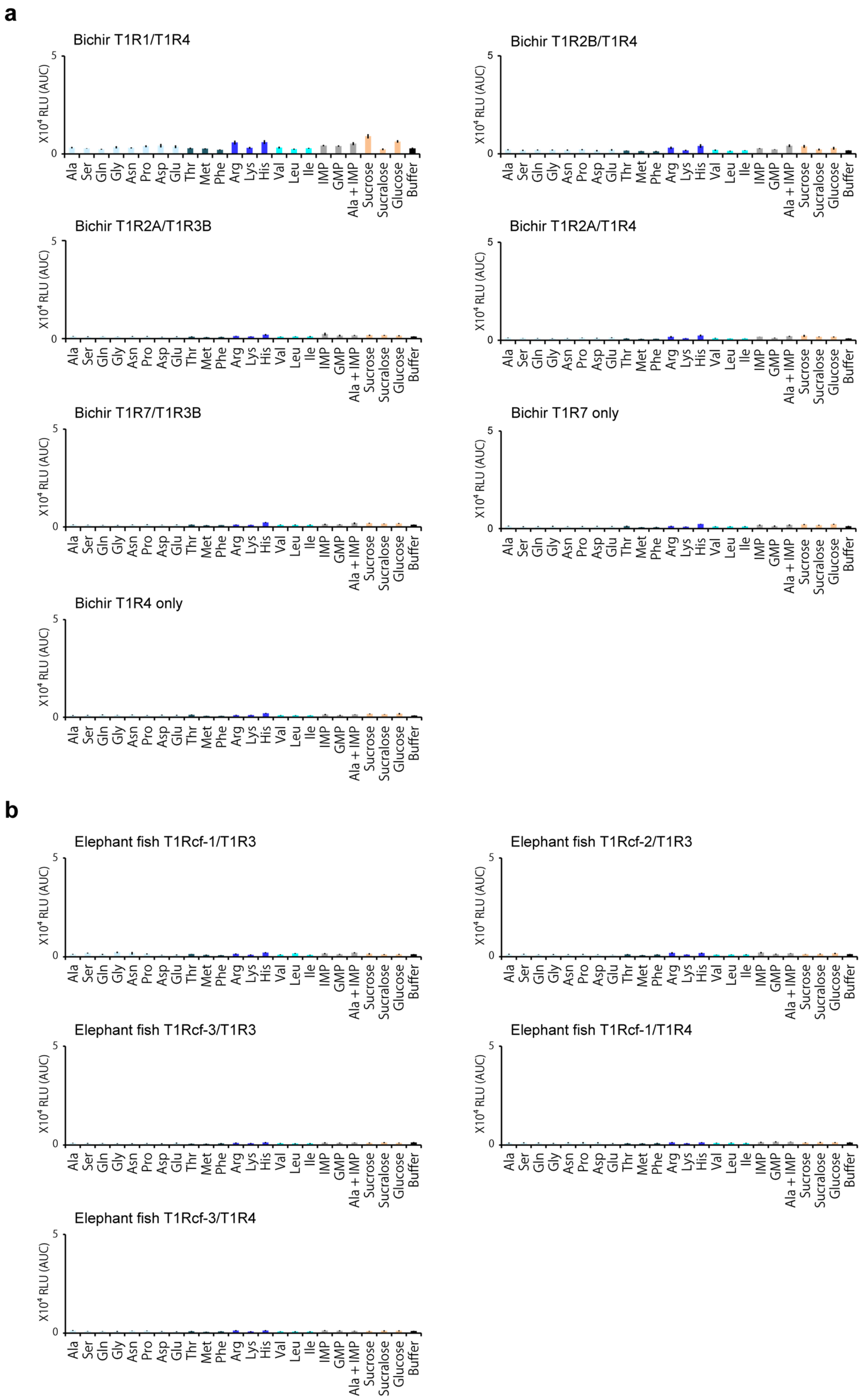


**Fig. S5. No significant response of various combinations of T1Rs.**

**a**, Five T1R combinations, T1R4-only, and T1R7-only from the bichir were coexpressed in HEK293T cells, and their responses to the 17 amino acids (50 mM), nucleic acids (10 mM), sugars and sucralose (100 mM) were tested. Values represent the mean ± standard error of six independent experiments performed with duplicate samples. **b**, Same as (**a**) except using five T1R combinations from the elephant fish.

**Table S1. Genome and transcriptome data used in this study.**

| Species | Data version and accession | Reference |
|---|---|---|
| Human (*Homo sapiens*) | GRCh38.p13 (NCBI: GCF_000001405.39) | 1 |
| Mouse (*Mus musculus*) | GRCm38.p6 (NCBI: GCF_000001635.26) | 2 |
| Chicken (*Gallus gallus*) | GRCg6a (NCBI: GCF_000002315.6) | 3 |
| Green anole (*Anolis carolinensis*) | AnoCar2.0 (NCBI: GCF_000090745.1) | 4 |
| Central bearded dragon (*Pogona vitticeps*) | pvi1.1 (NCBI: GCF_900067755.1) | 5 |
| Japanese gecko (*Gekko japonicas*) | Gekko_japonicus_V1.1 (NCBI: GCF_001447785.1) | 6 |
| Tibetan frog (*Nanorana parkeri*) | ASM93562v1 (NCBI: GCF_000935625.1) | 7 |
| Axolotl (*Ambystoma mexicanum*) | AmbMex13_14.1.0 (NCBI: GCA_001455525.1) | 8 |
| | RNA-seq (NCBI: PRJEB22921) | 9 |
| | RNA-seq (NCBI: PRJNA312389) | 10 |
| | RNA-seq (NCBI: PRJNA354434) | 11 |
| Two-lined caecilian (*Rhinatrema bivittatum*) | aRhiBiv1.1 (NCBI: GCF_901001135.1) | 12 |
| West African lungfish (*Protopterus annectens*) | National Genomic Data Center : GWHANVS00000000 | 13 |
| Australian lungfish (*Neoceratodus forsteri*) | neoFor_v3 (NCBI: GCA_016271365.1) | 14 |
| Coelacanth (*Latimeria chalumnae*) | LatCha1 (NCBI: GCF_000225785.1) | 15, 16 |
| Fugu (*Takifugu rubripes*) | FUGU5 (NCBI: GCF_000180615.1) | 17 |
| Zebrafish (*Danio rerio*) | GRCz11 (NCBI: GCF_000002035.6) | 18 |
| Spotted gar (*Lepisosteus oculatus*) | LepOcu1 (NCBI: GCF_000242695.1) | 19 |
| Bowfin (*Amia calva*) | AmiCal1 (NCBI: GCA_017591415.1) | 20 |
| Bichir (*Polypterus senegalus*) | AS1683550v1 (NCBI: GCF_016835505.1) | 20 |
| | polypterus_v0.62 | Okabe et al. (unpublished) |
| Elephant shark (*Callorhinchus milii*) | Callorhinchus_milii-6.1.3 (NCBI: GCF_000165045.1) | 21 |
| Whale shark (*Rhincodon typus*) | ASM164234v2 (NCBI: GCF_001642345.1) | 22 |
| Cloudy catshark (*Scyliorhinus torazame*) | Storazame_v1.0 (NCBI: GCA_003427355.1) | 23 |
| Brownbanded bamboo shark (*Chiloscyllium punctatum*) | Cpunctatum_v1.0 (NCBI: GCA_003427335.1) | 23 |

**Supplementary References**


1 Lander, E. S. *et al.* Initial sequencing and analysis of the human genome. *Nature* **409**, 860-921, doi:10.1038/35057062 (2001).

2 Mouse Genome Sequencing, C. *et al.* Initial sequencing and comparative analysis of the mouse genome. *Nature* **420**, 520-562, doi:10.1038/nature01262 (2002).

3 International Chicken Genome Sequencing, C. Sequence and comparative analysis of the chicken genome provide unique perspectives on vertebrate evolution. *Nature* **432**, 695-716, doi:10.1038/nature03154 (2004).

4 Alfoldi, J. *et al.* The genome of the green anole lizard and a comparative analysis with birds and mammals. *Nature* **477**, 587-591, doi:10.1038/nature10390 (2011).

5 Georges, A. *et al.* High-coverage sequencing and annotated assembly of the genome of the Australian dragon lizard Pogona vitticeps. *Gigascience* **4**, 45, doi:10.1186/s13742-015-0085-2 (2015).

6 Liu, Y. *et al.* Gekko japonicus genome reveals evolution of adhesive toe pads and tail regeneration. *Nat Commun* **6**, 10033, doi:10.1038/ncomms10033 (2015).

7 Sun, Y. B. *et al.* Whole-genome sequence of the Tibetan frog Nanorana parkeri and the comparative evolution of tetrapod genomes. *Proc Natl Acad Sci U S A* **112**, E1257-1262, doi:10.1073/pnas.1501764112 (2015).

8 Nowoshilow, S. *et al.* The axolotl genome and the evolution of key tissue formation regulators. *Nature* **554**, 50-55, doi:10.1038/nature25458 (2018).

9 Thampi, P., Liu, J., Zeng, Z. & MacLeod, J. N. Changes in the appendicular skeleton during metamorphosis in the axolotl salamander (Ambystoma mexicanum). *J Anat* **233**, 468-477, doi:10.1111/joa.12846 (2018).

10 Jiang, P. *et al.* Analysis of embryonic development in the unsequenced axolotl: Waves of transcriptomic upheaval and stability. *Dev Biol* **426**, 143-154, doi:10.1016/j.ydbio.2016.05.024 (2017).

11 Caballero-Perez, J. *et al.* Transcriptional landscapes of Axolotl (Ambystoma mexicanum). *Dev Biol* **433**, 227-239, doi:10.1016/j.ydbio.2017.08.022 (2018).

12 Rhie, A. *et al.* Towards complete and error-free genome assemblies of all vertebrate species. *Nature* **592**, 737-746, doi:10.1038/s41586-021-03451-0 (2021).

13 Wang, K. *et al.* African lungfish genome sheds light on the vertebrate water-to-land transition. *Cell* **184**, 1362-1376 e1318, doi:10.1016/j.cell.2021.01.047 (2021).

14 Meyer, A. *et al.* Giant lungfish genome elucidates the conquest of land by vertebrates. *Nature* **590**, 284-289, doi:10.1038/s41586-021-03198-8 (2021).

15 Amemiya, C. T. *et al.* The African coelacanth genome provides insights into tetrapod evolution. *Nature* **496**, 311-316, doi:10.1038/nature12027 (2013).

16 Nikaido, M. *et al.* Coelacanth genomes reveal signatures for evolutionary transition from water to land. *Genome Res* **23**, 1740-1748, doi:10.1101/gr.158105.113 (2013).

17 Aparicio, S. *et al.* Whole-genome shotgun assembly and analysis of the genome of Fugu rubripes. *Science* **297**, 1301-1310, doi:10.1126/science.1072104 (2002).

18 Howe, K. *et al.* The zebrafish reference genome sequence and its relationship to the human genome. *Nature* **496**, 498-503, doi:10.1038/nature12111 (2013).

19 Braasch, I. *et al.* The spotted gar genome illuminates vertebrate evolution and facilitates human-teleost comparisons. *Nat Genet* **48**, 427-437, doi:10.1038/ng.3526 (2016).

20 Bi, X. *et al.* Tracing the genetic footprints of vertebrate landing in non-teleost ray-finned fishes. *Cell* **184**, 1377-1391 e1314, doi:10.1016/j.cell.2021.01.046 (2021).

21 Venkatesh, B. *et al.* Elephant shark genome provides unique insights into gnathostome evolution. *Nature* **505**, 174-179, doi:10.1038/nature12826 (2014).

22 Read, T. D. *et al.* Draft sequencing and assembly of the genome of the world's largest fish, the whale shark: Rhincodon typus Smith 1828. *BMC Genomics* **18**, 532, doi:10.1186/s12864-017-3926-9 (2017).

23 Hara, Y. *et al.* Shark genomes provide insights into elasmobranch evolution and the origin of vertebrates. *Nat Ecol Evol* **2**, 1761-1771, doi:10.1038/s41559-018-0673-5 (2018).
